# GAN-based anomaly detection in multi-modal MRI images

**DOI:** 10.1101/2020.07.10.197087

**Authors:** Sean Benson, Regina Beets-Tan

**Affiliations:** Department of Radiology, the Netherlands Cancer Institute, Amsterdam, The Netherlands; GROW School for Oncology and Developmental Biology, Maastricht University Medical Center, Maastricht, The Netherlands

## Abstract

Generative adversarial networks (GANs) are known to be a powerful tool in order to correct image aberrations, and even predict entirely synthetic images. We describe and demonstrate a method to use GANs trained from multi-modal magnetic resonance images as a 3-channel input. The training of the generative network was performed using only healthy images together with pseudo-random irregular masks. The dataset consisted of just 20 people. The resulting model was then used to detect anomalies real patient images in which the anomaly was a tumour. The search was performed using no prior knowledge of the tumour location, if indeed a tumour was present. Resulting accuracies are observed to vary significantly on the size of the anomaly. The area under the receiver operator characteristic curve is observed to be greater than 0.75 for anomaly sizes greater than 4 cm^2^.

## Introduction

Generative adversarial networks (GANs) consist of a generator network designed to produce simulated data and a discriminator network designed to discriminate real from synthetic images. The generator and discriminator networks are trained together in a zero sum game, with the ideal result that the generator network is able to produce entirely realistic images. Such networks have been proven to be extremely versatile, with usage examples varying from image correction [1], and background removal [2], to the generation of new works of entirely new images from a text description [3]. The training of such networks from scratch often requires 10s of thousands to more than 100,000 images to achieve acceptable performance. In the context of medical imaging, GANs have already been used to correct distortions in images, standardise images from different scanners [4], and even generate entire synthetic 3D brain volumes [5]. When training GANs it is common to use a network that has been pre-trained with a large dataset, and subsequently perform a fine-tuning on the available dataset. This has led to reliable complete images being generated from masked magnetic resonance images (MRI) [6]. A review of 150 publications detailing the use case of GANs for medical imaging is given in Ref. [7].

In this work, we detail the training of a GAN using the method of partial convolutions [8] with healthy MRI images of the brain from just 20 patients. The partial convolution layer involves performing the convolution operation only on regions of the image in which a mask is present to some degree, where the mask has been initialised to block random parts of training images for the GAN to complete. Crucially, this also means the mask is updated based on whether the convolution was able to condition its output on at least one valid input value. Ideally, successive applications of the method mean that the mask eventually disappears.

The model, once trained, was then used to scan a separate set of images via a custom search algorithm, in which the location of a tumour and indeed whether a tumour is present is not provided to the model. The ability of the algorithm to identify anomalous areas of the scan arising from the presence of a tumour is then assessed.

## Method

A dataset consisting of 20 patients from The Cancer Imaging Archive was used [9, 10]. For each patient MRI scans from two different time points were performed, using a variety of modalities. The two time points corresponded to within 90 days of chemo-radiotherapy treatment completion and at a clinically determined point of progression. The MRI modalities contained in the dataset consisted of T1-weighted (pre- and post-contrast agent), T2-weighted, fluid-attenuated inversion recovery (FLAIR), apparent diffusion coefficient (ADC), normalised cerebral blood flow, normalised relative cerebral blood volume, and standardized relative cerebral blood volume. While the manufacturer and model was not always the same, the dataset was found to be consistent in terms of pixel spacing (0.6875mm) and axial slice spacing (6.5mm). Multihance was also found to be consistently used as the contrast agent.

In order to ensure that only the brain volume was processed as input to the neural network, a brain mask was generated from NiLearn [11] on the basis of the T1-weighted MRI images (post contrast). In the event that NiLearn was unable to provide a mask for a patient MRI scan, the data point was skipped. A three-channel image was then created using the FLAIR (red-channel), ADC (green-channel), and dT1 (blue-channel) images, where dT1 is the difference between the T1-weighted images pre- and post-contrast. The provided tumour mask was then used to separate healthy axial slices and slices in which a tumour was present. Example images in which a tumour was present are shown in Fig. 1, after application of the brain volume mask. The total dataset of 308 images was then obtained which were split into training (n=109), test (n=104), and validation (n=95) on the basis of axial slice number modulo 3.

**Fig 1.**
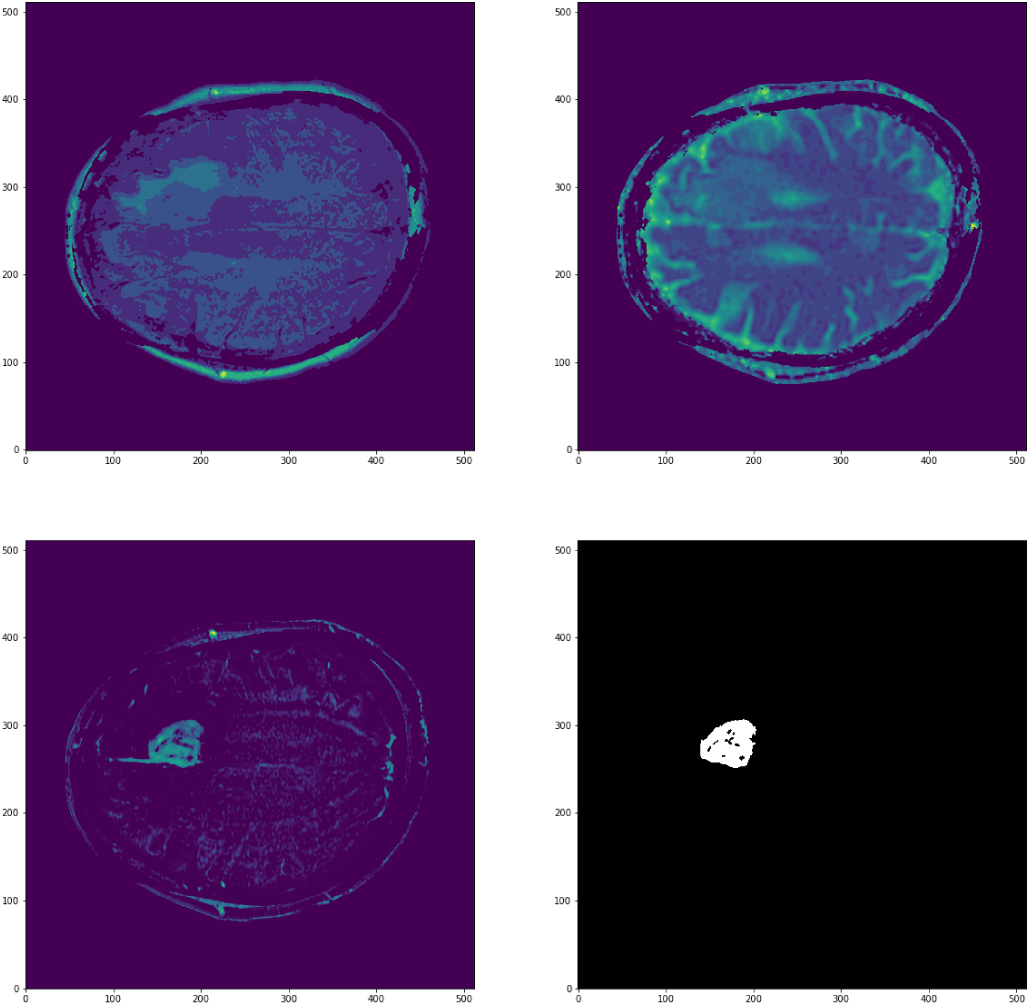
Example axial slice from the TCIA dataset [10] of the FLAIR (top-left), ADC (top-right), and dT1 (bottom-left) images, together with the provided tumour mask (bottom-right).

The GAN model was based on the Keras implementation [12] of the partial convolution (PConv) method of Ref. [8]. A starting point for the training was taken from the pre-existing training to the ImageNet dataset [12]. In total 500 epochs were performed with a learning rate of 0.0002, after which improvement became negligible. Training was performed using the same irregular mask method of Ref. [12], in which masks are generated from multiple random geometric shapes and lines. Example masked images are shown in Fig. 2.

**Fig 2.**
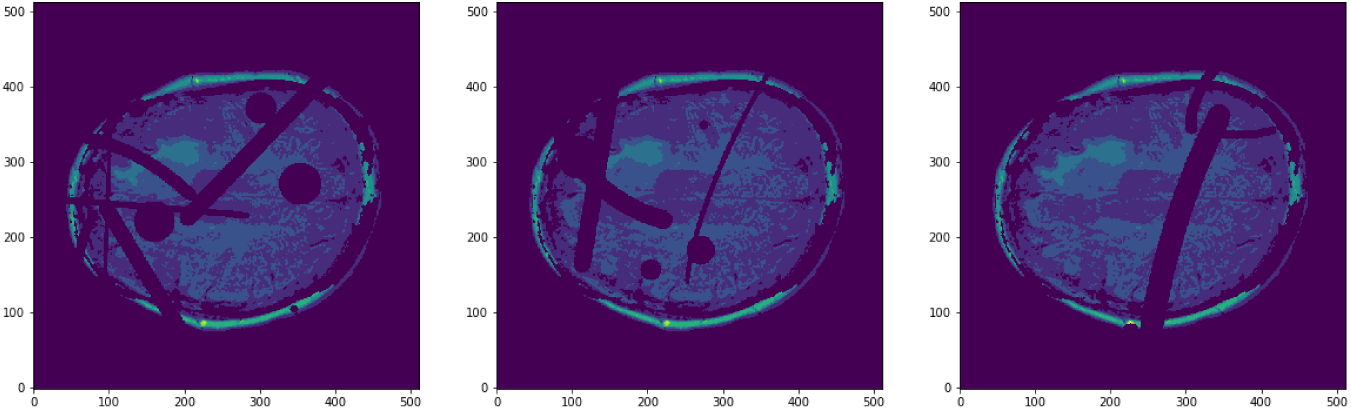
Examples of irregular masks made from lines of varying thickness and circles of varying size applied to the FLAIR channel.

In order to detect anomalies, a sequence of scans is made through the image using rectangular masks of varying size. The model is repeatedly evaluated moving the mask through the image until the entire image has been probed. The presence of an anomaly is inferred through the value of *a*_*k*_, defined as

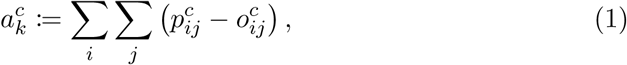

where *k* is the mask width of the scan, *c* is the image channel (dT1, FLAIR or ADC), *i, j* are pixels in the mask region, *p*_*ij*_ is the value of the pixel in the predicted image and *o*_*ij*_ is the value of the pixel in the original image. An anomaly is said to be detected if 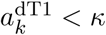, where *κ* is a configurable threshold. This then indicates that increased signal intensity is seen in the image above that expected by the neural network. The results of scans with *k* = 50, *k* = 60, and *k* = 70 are combined.

## Results and discussion

A comparison of the predicted images with the original images from the test dataset using the trained model is shown in Fig. 3. Good agreement can be seen in the dT1 channel whereas a noticeable reduction in signal intensity is present for the case of the predicted FLAIR and ADC channels.

**Fig 3.**
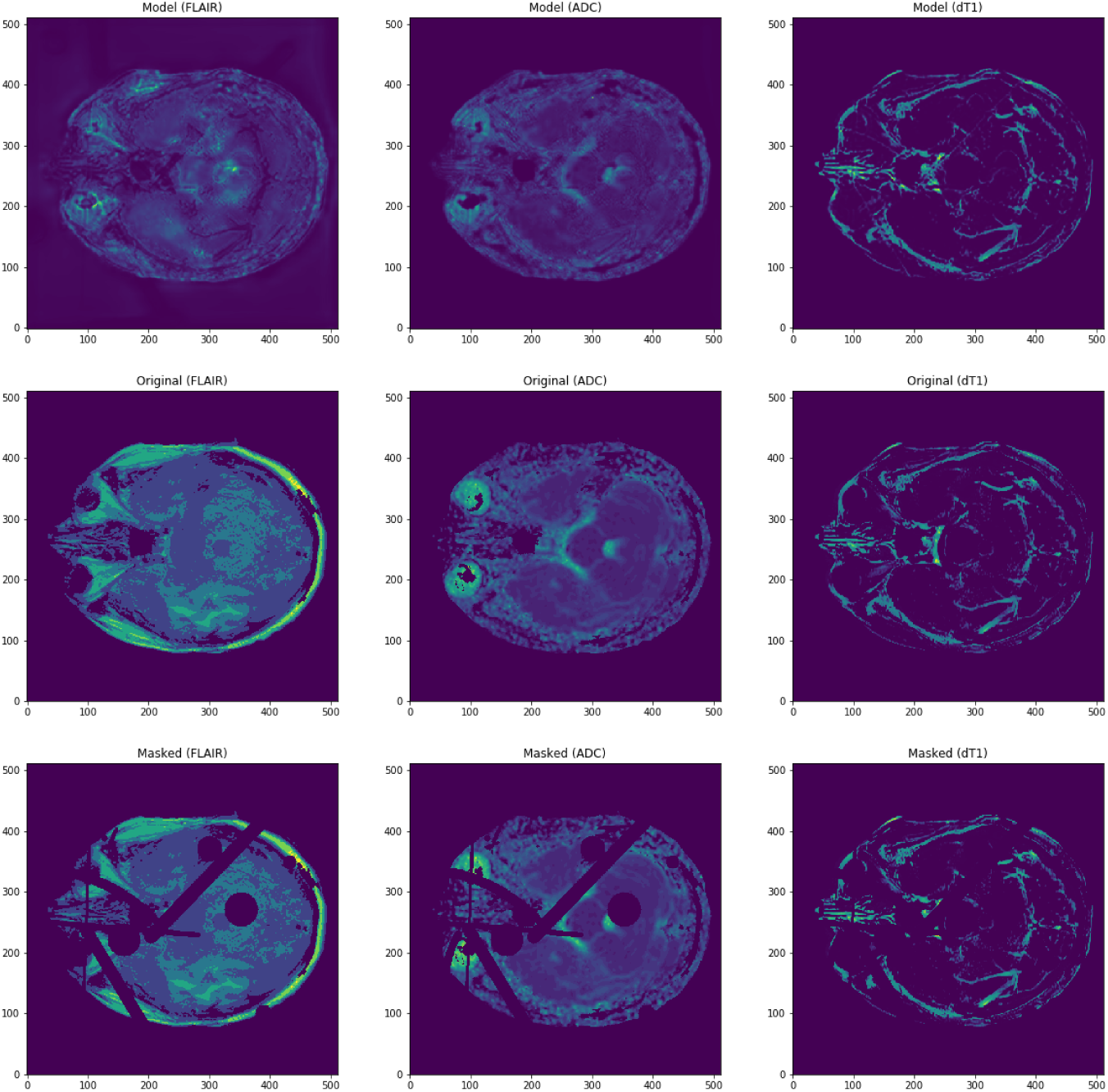
Comparison of model predicted images against the ground truth for an image in the test dataset.

The results of the anomaly scan method described earlier are evaluated using the axial slice shown in Fig. 1. This is compared against the original image in Fig. 4. The anomalous area is seen to show good agreement with the area of the tumour mask in Fig. 1.

**Fig 4.**
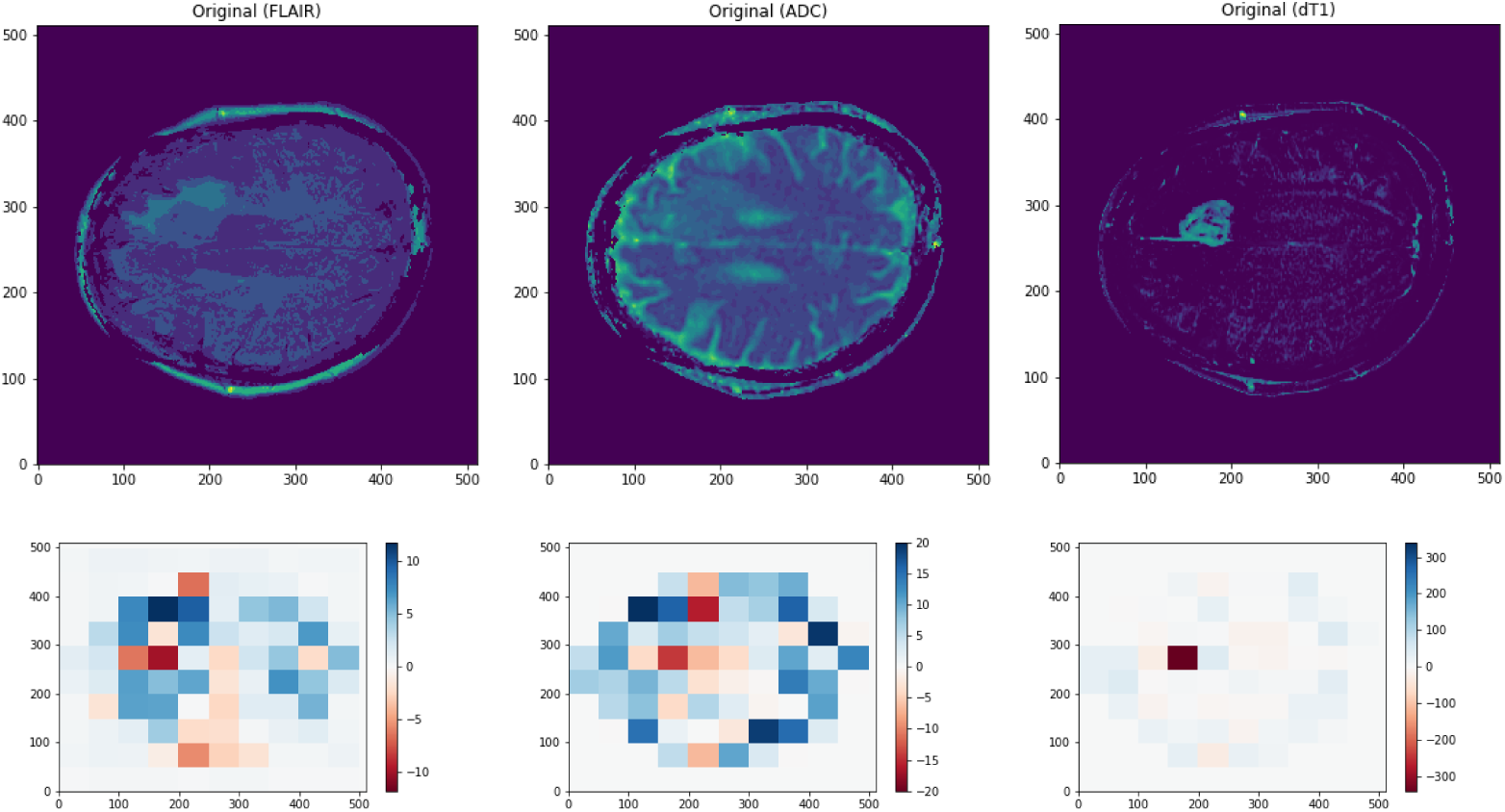
Comparison of model scan results against an example image with a tumour present.

The performance of the simple search anomaly detection method was evaluated using the image slices in which a tumour was determined to be present from the provided tumour mask. This therefore represents a dataset that is completely independent from the training, test, and validation samples created from the healthy images. An anomaly is only considered successfully detected if there is overlap between the anomalous region and the provided tumour mask. The results of the simple anomaly scan method described earlier are shown in Fig. 5 as a function of tumour area, which is determined using the pixel spacing of 0.6875 mm found from the dicom image header. The receiver operator characteristic curves, shown in Fig. 5 were generated varying the value of *κ* in the range (−500, 40).

**Fig 5.**
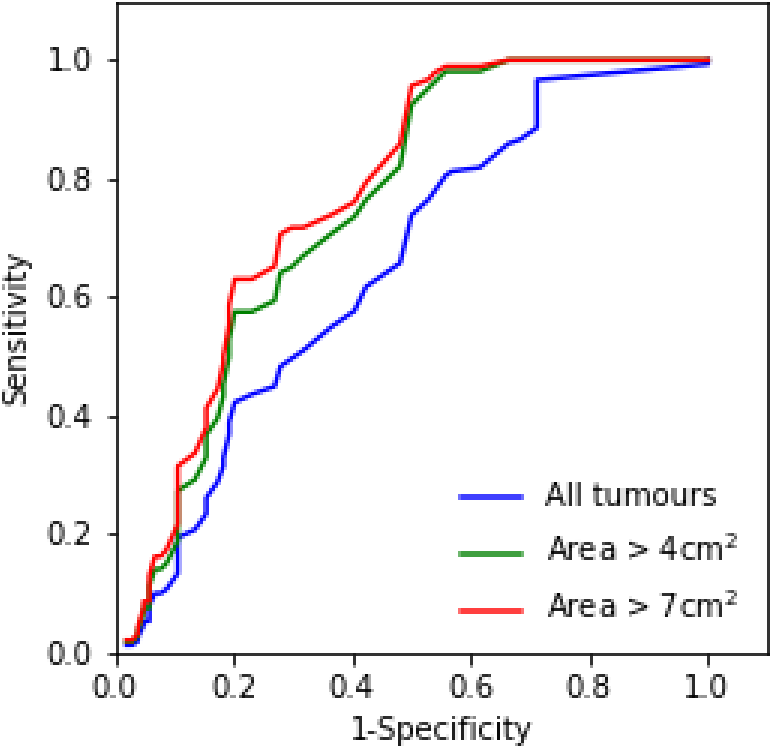
Receiver operator characteristic curves for different anomaly sizes.

The values of the area under the curve were determined to be 0.65, 0.75, and 0.77, cross section area greater than 0, 4, and 7 cm^2^, respectively. As would be naively expected, performance is seen to decrease as a function of tumour area, though consistently remains above the false-positive rate.

The ability of the network to discern anomalies is limited by the amount of training data, thus improved performance is expected for smaller tumour sizes with increasing training set size. It will be important to test this assumption with larger training datasets. An important feature of the method is that no images of cancerous tissue were used in the training (under the assumption that the provided tumour masks were accurate), and therefore no labels of whether tumour tissue is present are relevant. The use of a simple grid search in order to detect anomalies can be optimised further and is included here to demonstrate the capabilities. More advanced methods exist and will be useful in subsequent analyses on larger datasets (the interested reader is directed to a recent review in Ref. [13]).

It is important to note that while MRI images are used in this example, the method should be applicable to other modalities. An area under active research for artificial intelligence is gastrointestinal endoscopy, where AI can be used in the context of lesion detection and possible cancer regrowth [14]. The potentially large dataset sizes and variable camera position could potentially make the approach powerful.

## Conclusion

We have demonstrated a method by which a generative adversarial network may be used to detect tumours with only healthy data used in training. Encouraging performance is seen from a relatively small dataset of 20 patients together with a rule-based anomaly detection algorithm. Given the large quantities of available healthy data, subsequent studies using larger datasets and more advanced anomaly detection algorithms will be interesting to see the performance limits, especially in terms of anomaly size.

## References

1. S. Tripathi, Z. C. Lipton, and T. Q. Nguyen, “Correction by Projection: Denoising Images with Generative Adversarial Networks,” arXiv e-prints, Mar. 2018, 1803.04477.

2. T. Bouwmans, S. Javed, M. Sultana, and S. K. Jung, “Deep Neural Network Concepts for Background Subtraction: A Systematic Review and Comparative Evaluation,” arXiv e-prints, Nov. 2018, 1811.05255.

3. H. Zhang, T. Xu, H. Li, S. Zhang, X. Huang, X. Wang, and D. N. Metaxas, “Stackgan: Text to photo-realistic image synthesis with stacked generative adversarial networks,” arXiv e-prints, 2016, 1612.03242.

4. K. Armanious, C. Jiang, M. Fischer, T. Küstner, K. Nikolaou, S. Gatidis, and B. Yang, “MedGAN: Medical Image Translation using GANs,” arXiv e-prints, June 2018, 1806.06397.

5. G. Kwon, C. Han, and D.-s. Kim, “Generation of 3d brain mri using autoencoding generative adversarial networks,” in Medical Image Computing and Computer Assisted Intervention – MICCAI 2019 (D. Shen, T. Liu, T. M. Peters, L. H. Staib, C. Essert, S. Zhou, P.-T. Yap, and A. Khan, eds.), (Cham), pp. 118–126, Springer International Publishing, 2019.

6. K. Armanious, V. Kumar, S. Abdulatif, T. Hepp, S. Gatidis, and B. Yang, “ipA-MedGAN: Inpainting of Arbitrary Regions in Medical Imaging,” arXiv e-prints, Oct. 2019, 1910.09230.

7. X. Yi, E. Walia, and P. Babyn, “Generative adversarial network in medical imaging: A review,” Medical Image Analysis, vol. 58, p. 101552, 2019.

8. G. Liu, F. A. Reda, K. J. Shih, T. Wang, A. Tao, and B. Catanzaro, “Image inpainting for irregular holes using partial convolutions,” CoRR, vol. abs/1804.07723, 2018, 1804.07723.

9. K. Clark, B. Vendt, K. Smith, J. Freymann, J. Kirby, P. Koppel, S. Moore, S. Phillips, D. Maffitt, M. Pringle, L. Tarbox, and F. Prior, “The cancer imaging archive (TCIA): Maintaining and operating a public information repository,” Journal of Digital Imaging, vol. 26, pp. 1045–1057, July 2013.

10. K. Schmainda and M. Prah, “Data from brain-tumor-progression,” 2019.

11. A. Abraham, F. Pedregosa, M. Eickenberg, P. Gervais, A. Mueller, J. Kossaifi, A. Gramfort, B. Thirion, and G. Varoquaux, “Machine learning for neuroimaging with scikit-learn,” Frontiers in Neuroinformatics, vol. 8, 2014.

12. M. Gruber, “Partial convolutions for image inpainting using keras.” https://github.com/MathiasGruber/PConv-Keras, 2019.

13. M. Goldstein and S. Uchida, “A comparative evaluation of unsupervised anomaly detection algorithms for multivariate data,” PLOS ONE, vol. 11, p. e0152173, Apr. 2016.

14. A. P. Abadir, M. F. Ali, W. Karnes, and J. B. Samarasena, “Artificial intelligence in gastrointestinal endoscopy,” Clinical Endoscopy, vol. 53, pp. 132–141, Mar. 2020.

